# Low Transcriptional Complexity Cells Represent a Conserved, Aging-Relevant Maintenance State Overlooked by Single-Cell Transcriptomics

**DOI:** 10.64898/2026.02.16.706190

**Authors:** Alexander Bontempo, Prabhu Mathiyalagan

## Abstract

Whether mature tissues harbor transcriptionally quiet yet biologically functional cellular reservoirs remains largely unexplored. Here, using publicly available single-cell and single-nucleus transcriptomic datasets, we identify a previously unrecognized cellular state within fully differentiated human cell lineages characterized by low transcriptomic complexity (<1000 genes per cell) but preserved lineage identity and coherent functional gene expression. These “low-transcriptional” (low-T) states, often excluded by standard single-cell quality control thresholds or subsumed within major populations, are widespread across major organs including heart, brain, lung, and immune system, comprising substantial fractions of nearly all mature cell types. Despite reduced transcript abundance, low-T cells exhibit organized molecular programs distinct from high-T counterparts enriched in pathways related to cellular maintenance, metabolic resilience, survival, and aging, while lacking stress, apoptosis, senescence, or inflammation signatures. Low-T programs are conserved across mature cell lineages within organs but remain tissue-specific, revealing a hidden axis of cellular organization orthogonal to cell identity. Their abundance declines with age in brain and immune tissues, linking this state to organismal aging. Together, our findings uncover a transcriptionally quiescent yet functionally mature cellular state conserved across tissues. This previously overlooked population represents a biologically meaningful reservoir with implications for tissue maintenance, longevity, regenerative biology, and potential pharmacological aging interventions, and challenges conventional interpretations of transcriptional sparsity in single-cell genomics.

## Introduction

Single-cell and single-nucleus RNA sequencing (scRNA-seq/snRNA-seq) have transformed our understanding of tissue organization by enabling unbiased profiling of gene expression across millions of individual cells^1,2,3^. These technologies have revealed previously unrecognized cell types, transient states, and lineage hierarchies, reshaping models of development, homeostasis, and disease^4,5,6^. Standard analytical pipelines, however, rely heavily on quality-control (QC) thresholds that prioritize cells with high gene counts and unique molecular identifiers (UMIs), routinely excluding cells with low transcriptional complexity. Many studies apply thresholds excluding cells with fewer than 200-1500 detected genes per cell^7,8,9^. While such filters reduce technical artifacts and ambient RNA contamination, they also embed the assumption that transcriptionally sparse cells are low quality, dying, or biologically uninformative. Although some studies question these QC practices, the assumption that low-transcript cells lack biological meaning remains largely untested^10,11^.

Mature, terminally differentiated cells, including cardiomyocytes, neurons, endothelial cells, fibroblasts, and immune cells, can be long-lived, metabolically quiescent, and transcriptionally stable^12^. These properties suggest that biologically meaningful cellular states may exist with intrinsically low transcriptional output^13^. Yet, such cells are systematically underrepresented in scRNA-seq studies due to aggressive filtering. Even among QC-passed cells, low-transcriptional-complexity cells are often subsumed into major lineages based solely on marker expression, without separate consideration of their transcriptional state. An approach explicitly stratifying cells by low versus high transcriptional complexity within mature lineages has not been applied, potentially obscuring important aspects of tissue organization, functional heterogeneity, and aging biology. We term these cells “low-T” cells, in contrast to transcriptionally complex “high-T” cells, and hypothesize that low-T states represent fully differentiated, lineage-committed cells maintained in a low-transcriptional-output state for functional or regulatory reasons. Such states may reflect quiescence, metabolic optimization, epigenetic stabilization, or adaptive responses to long-term tissue stress. Unlike progenitor or stem cells, which are transcriptionally active and plastic, low-T cells are predicted to be stable, mature, and lineage-specific.

Evidence from spatial transcriptomics and single-cell doublet analyses highlights that signals traditionally dismissed as technical artifacts can encode meaningful biology. Unassigned transcripts in spatial datasets reflect synaptic organization, cellular projections, and extracellular RNA localization^14^, and cells excluded as doublets in cancer scRNA-seq reveal biologically significant cellular interactions^15^. These observations illustrate a broader conceptual gap: signals falling outside conventional analytical frameworks may encode overlooked biological structure. By analogy, low-T cells, routinely filtered or overlooked in scRNA-seq and snRNA-seq datasets, may constitute a biologically meaningful state that has remained largely unexplored.

Aging provides a compelling biological context for low-T cells. Tissue aging is associated with declining regenerative capacity, altered cellular homeostasis, and accumulation of long-lived, differentiated cells with reduced plasticity^16,17^. Aging is characterized by progressive alterations in proteostasis, mitochondrial function, metabolic regulation, transcriptional fidelity, and inflammatory tone^18,19,20,21,22,23^. While these processes have been extensively described at the bulk tissue level and through aggregate single-cell analyses, it remains unclear whether such aging-associated programs are uniformly distributed across cells or preferentially enriched within specific transcriptional states. Most transcriptomic studies of aging focus on high-quality, transcriptionally active cells, potentially missing states linked to long-term cellular maintenance, resilience, or functional decline^24,25^. If low-T cells represent a stable mature state, changes in their abundance, lineage specificity, or transcriptional programs across the lifespan could reveal previously unrecognized aspects of tissue composition, cellular life history, and age-associated functional decline.

Here, we systematically investigate low-T cells across multiple human tissues, including heart, brain, lung, and immune compartments, using large-scale scRNA-seq and snRNA-seq datasets that have passed conventional QC thresholds. We ask whether low-T cells occur at substantial frequencies within major mature lineages, whether their distribution is stochastic or lineage-selective, and whether they exhibit reproducible transcriptional programs distinct from high-T counterparts within the same annotated cell types. We also evaluate whether these programs are conserved across tissues and individuals and how low-T states change across the human lifespan.

Our results reveal that low-T cells constitute a substantial fraction of cells across nearly all examined tissues and lineages, frequently comprising 10-90% of cells within canonical mature cell types. Despite reduced transcript counts, these cells exhibit coherent and reproducible gene expression programs distinct from high-T cells, including transcriptional signatures conserved across cell types within organs yet remaining tissue-specific. Notably, the prevalence of low-T states declines with age in brain and immune compartments, indicating a dynamic relationship between transcriptional sparsity and tissue aging.

Together, these findings challenge the prevailing assumption that low-transcript cells are merely technical artifacts and support the existence of a biologically meaningful, lineage-specific low-transcriptional state within mature tissues. By systematically characterizing this hidden transcriptional compartment, our study expands the conceptual landscape of cellular states accessible through single-cell transcriptomics. We propose that low-T cells represent a previously overlooked axis of cellular organization, orthogonal to cell type and lineage, with implications for tissue maintenance, aging, and potential rejuvenation. More broadly, our work underscores the importance of revisiting long-standing analytical conventions in genomics, as biologically informative signals may reside precisely in regions of data historically discarded.

## Results

### Low-T cells are widespread across major cell lineages in human tissues

To assess whether low-T cells represent a significant component of human scRNA-seq/snRNA-seq datasets, we analyzed published datasets that had already passed conventional QC filters (minimum 200 genes per cell; Table 1). We processed snRNA-seq data from human heart, brain, and lung, and scRNA-seq data from immune cells, followed by cell annotation. Cells were then stratified into Low-T (200-1000 detected genes) and High-T (>1000 genes). Low-T cells were embedded within nearly all major canonical cell types across tissues, present in both sexes and across young and old individuals (**Figure 1**). They did not form a separate cluster nor exhibit random distribution; instead, they were interspersed within each major cell type. Quantifying the proportion of low-T versus high-T cells within each lineage revealed that low-T cells constitute a substantial fraction of every major mature cell type, ranging from 10-90% across heart, brain, immune cells, and lung (**Figure 2**). These findings indicate that low-T cells are pervasive, embedded within mature lineages, and not segregated as independent clusters across human organs.

**Figure 1.**
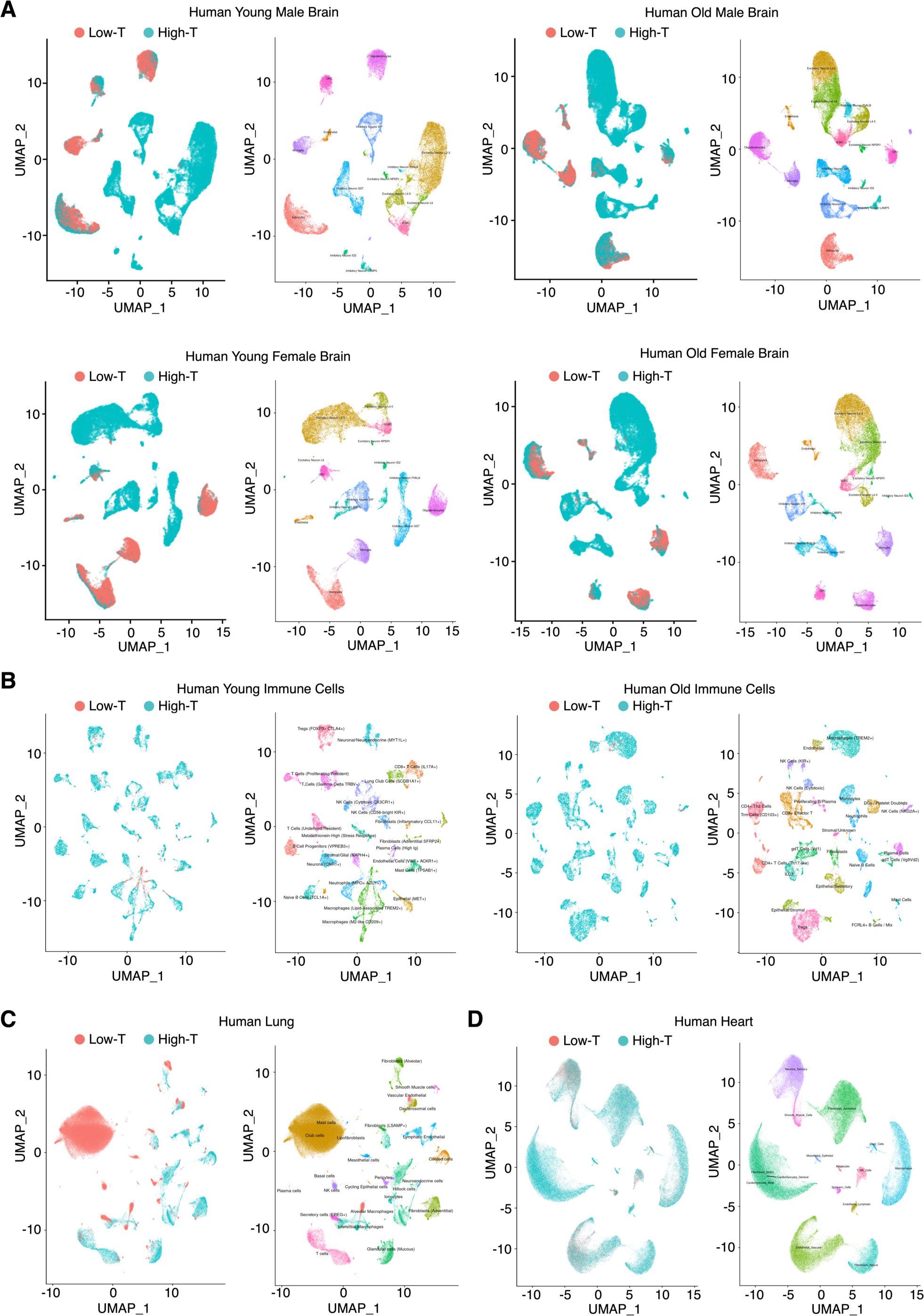
Stratification by transcriptional complexity reveals low-T cells embedded within canonical mature cell types across human tissues. (**A**) Human brain snRNA-seq datasets from young males (n = 6), old males (n = 6), young females (n = 3), and old females (n = 4). (**B**) Human immune cell scRNA-seq datasets from young (n = 1) and old (n = 1) donors. (**C**) Human lung snRNA-seq datasets (n = 4 donors). (**D**) Human heart snRNA-seq datasets (n = 25 donors). UMAPs are colored by transcriptional complexity state and annotated cell identity as indicated.

**Figure 2.**
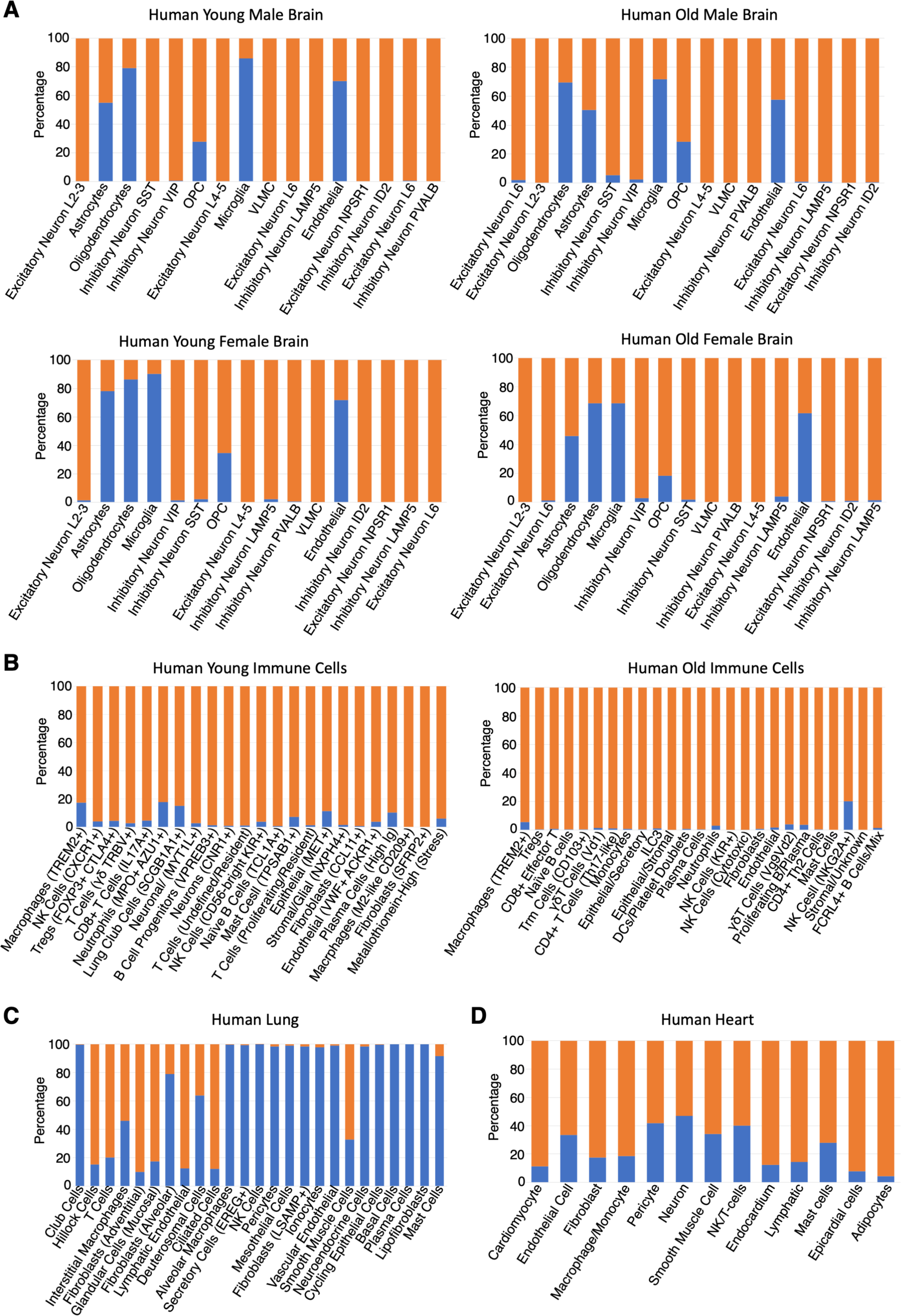
Proportion of low-T and high-T cells across annotated cell types in human tissues. Stacked bar plots show the proportion of low-T (<1,000 detected genes per cell) and high-T (≥1,000 genes per cell) cells within each annotated mature cell type. (**A**) Human brain snRNA-seq datasets from young males, old males, young females, and old females. (**B**) Human immune cell scRNA-seq datasets from young and old donors. (**C**) Human lung snRNA-seq datasets. (**D**) Human heart snRNA-seq datasets. Proportions were calculated within annotated cell types after quality control filtering.

### Low-T cells are transcriptionally distinct from high-T counterparts within the same lineage

We next asked whether low-T and high-T states within the same cell type display distinct transcriptional signatures. Differential gene expression analysis revealed numerous genes significantly upregulated in low-T cells compared to high-T cells across all tissues (**Figure 3, Table 2**). Similarly, low-T cells exhibited selective downregulation of other genes relative to high-T counterparts (**Figure 4, Table 3**). Importantly, low-T cells were not merely a subset of downregulated high-T cells. Instead, they displayed selective upregulation of genes reflecting qualitative changes in transcriptional programs specific to each cell type, indicating that low-T represents a distinct transcriptional state, embedded within the same annotated lineage.

**Figure 3.**
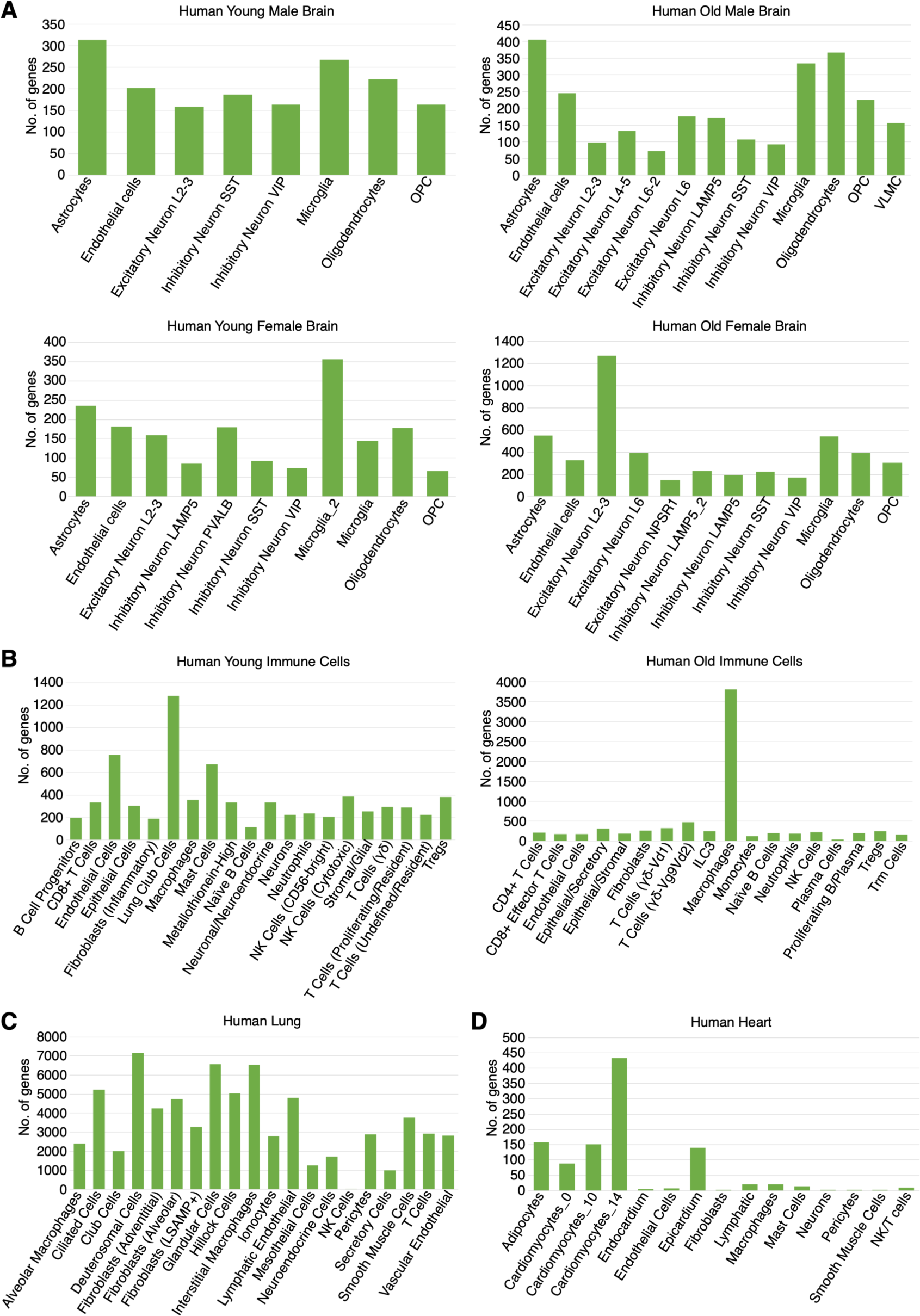
Number of genes upregulated in low-T cells relative to high-T cells across human tissues. (**A**) Human brain snRNA-seq datasets from young males, old males, young females, and old females. (**B**) Human immune cell scRNA-seq datasets from young and old donors. (**C**) Human lung snRNA-seq datasets. (**D**) Human heart snRNA-seq datasets. Differential expression analysis was performed within annotated cell types following quality control filtering. Genes were considered significantly upregulated at p < 0.05.

**Figure 4.**
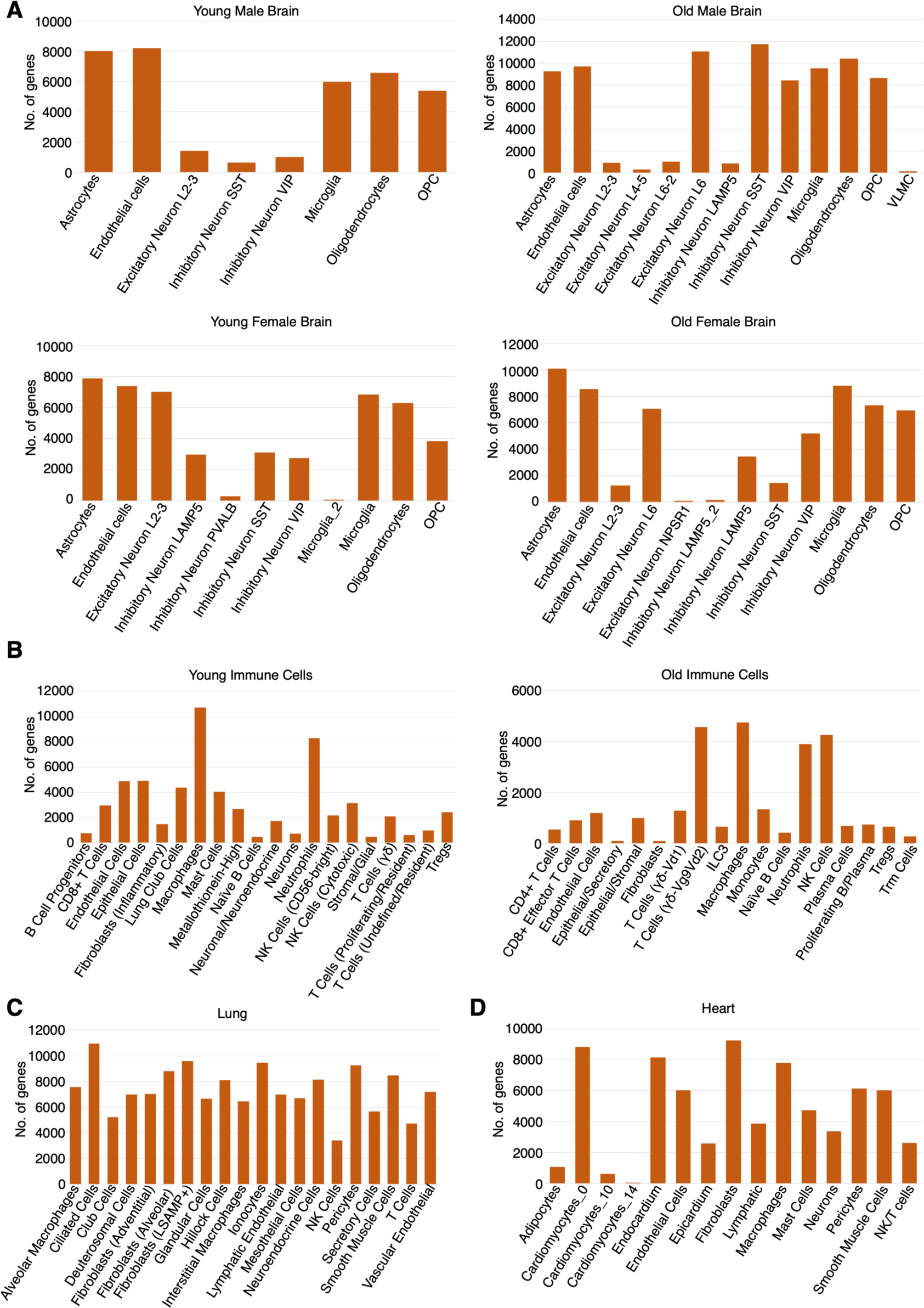
Number of genes downregulated in low-T cells relative to high-T cells across human tissues. (**A**) Human brain snRNA-seq datasets from young males, old males, young females, and old females. (**B**) Human immune cell scRNA-seq datasets from young and old donors. (**C**) Human lung snRNA-seq datasets. (**D**) Human heart snRNA-seq datasets. Differential expression analysis was performed within annotated cell types following quality control filtering. Genes were considered significantly upregulated at p < 0.05.

### Low-T cells share commonly upregulated genes across major cell types within each organ

We then examined whether these low-T-specific genes are conserved across mature lineages within the same organ. Intersecting upregulated genes across major cell types revealed substantial overlap within organs. In brain, 83, 125, 82, and 150 genes were commonly upregulated in low-T cells across mature cell types in young males, old males, young females, and old females, respectively (**Figure 5A, Table 4**). In immune cells, 91 genes were shared across NK cells, neutrophils, epithelial cells, and Tregs, 80 genes across macrophages, mast cells, endothelial, and neuroendocrine cells, and 66 genes common to all eight cell types (**Figure 5B, Table 4**). In lung, 1,571 genes were shared among macrophage, deuterosomal, hillock, and lymphatic endothelial cells, while 577 genes across T cells, smooth muscle, vascular endothelial, and pericytes, and 417 genes common to all eight cell types (**Figure 5C, Table 4**).

**Figure 5.**
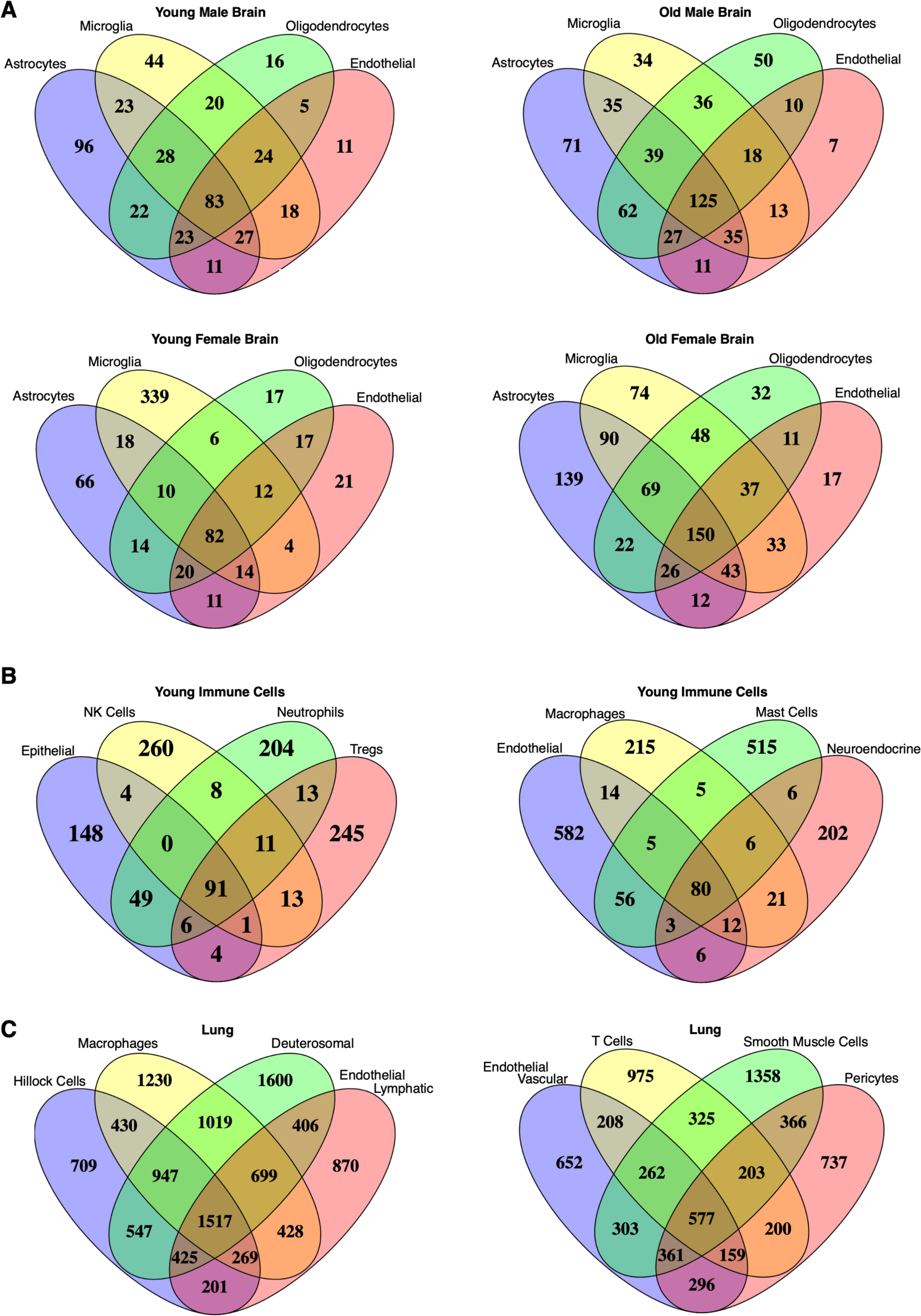
Overlap of genes upregulated in low-T cells across multiple mature cell types within each organ. Venn diagrams show the intersection of genes significantly upregulated in low-T cells relative to high-T cells across annotated mature cell types within each organ. (**A**) Upregulated genes in low-T cells from astrocytes, microglia, oligodendrocytes, and endothelial cells in human brain snRNA-seq datasets. (**B**) Upregulated genes in low-T cells from multiple mature cell types in young human immune cell scRNA-seq datasets. (**C**) Upregulated genes in low-T cells from multiple mature cell types in human lung snRNA-seq datasets.

Comparison of inter-individual gene programs in the brain showed 31 genes commonly upregulated in young males and females, 89 genes in old males and females, 63 genes in young vs old males, and 32 genes in young vs old females, with 18 genes consistently shared across all brain datasets (**Figure 6A, Table 5**). Cross-organ comparison revealed no genes shared across brain, immune, and lung, suggesting that low-T programs are conserved within organs but tissue-specific (**Figure 6B**). These results indicate that low-T cells exhibit organ-specific transcriptional programs that are also partially conserved across individuals.

**Figure 6.**
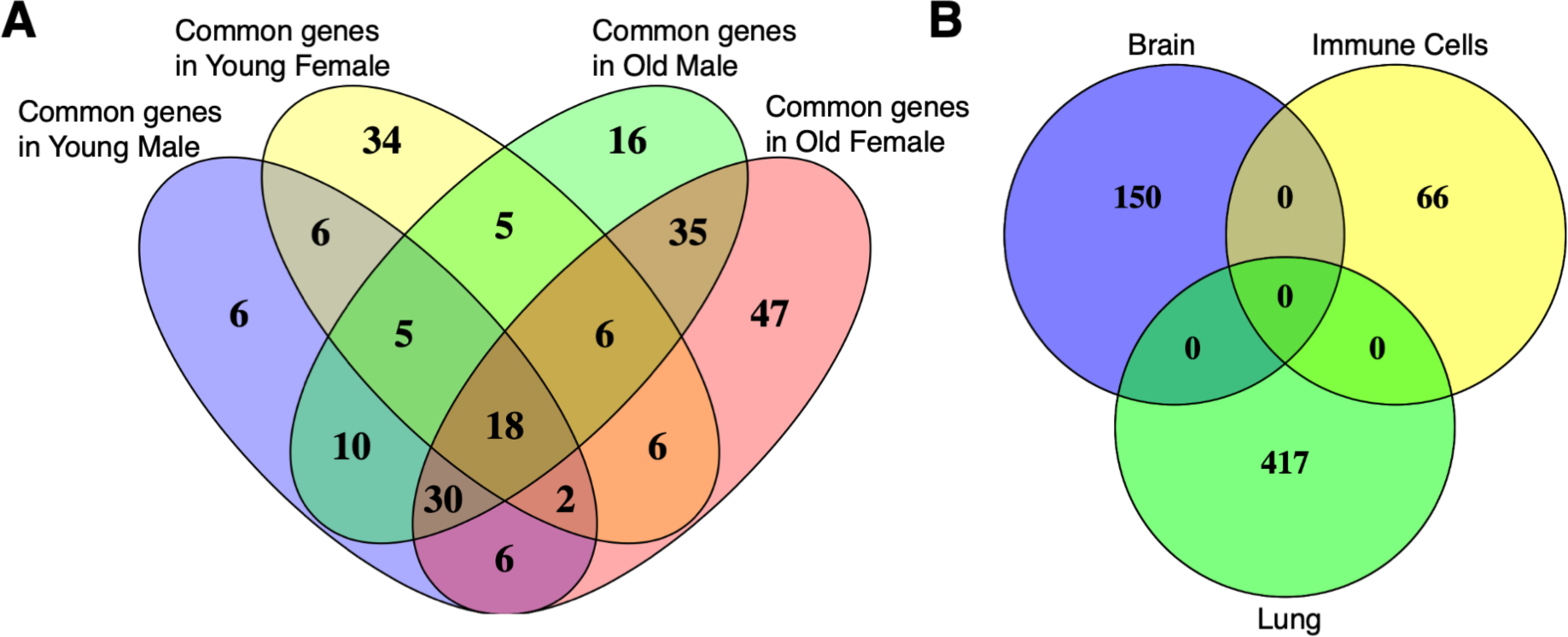
Comparison of genes upregulated in low-T cells across brain demographic groups and across tissues. (**A**) Genes upregulated in low-T cells from young males, old males, young females, and old females in human brain datasets were intersected to identify genes shared across all demographic groups. (**B**) Genes upregulated in low-T cells were intersected across tissues (brain, immune, and lung) to assess shared gene overlap.

### Low-T cells display coherent, biologically meaningful programs

We then sought to investigate whether the shared gene programs among low-T cells show biologically meaningful programs. To do this, we carried out pathway analyses. Pathway analysis revealed that the shared upregulated genes in low-T cells are functionally coherent rather than random. In brain, low-T cells show enrichment for synaptic and excitability pathways, including neuronal system signaling, chemical synaptic transmission, NMDA/AMPA receptor activation, ion channel transport, voltage-gated potassium channels, and long-term potentiation (**Figure 7A, Table 6**). Low-T neurons exhibit a compressed yet functionally specialized state, prioritizing energy-efficient synaptic maintenance.

**Figure 7.**
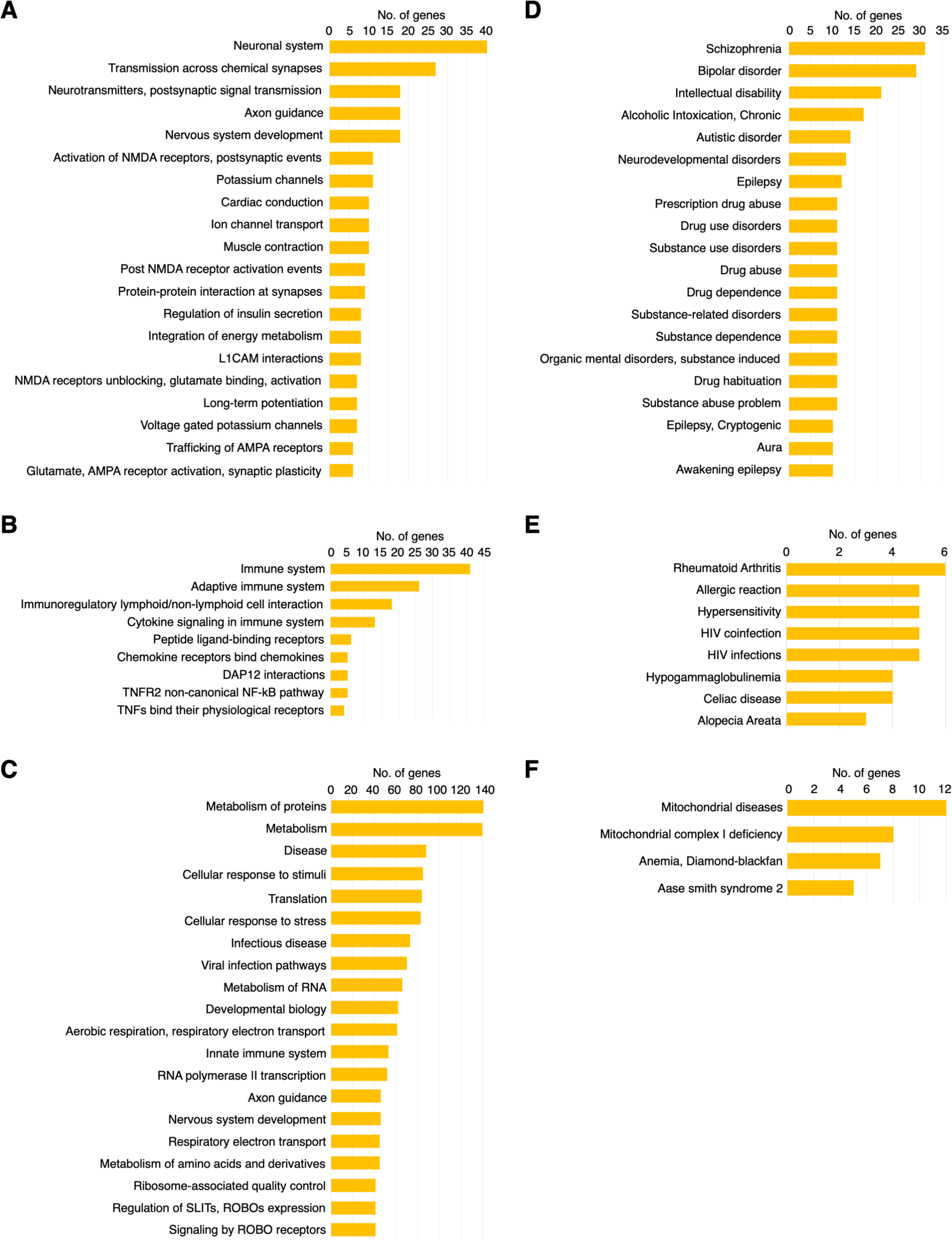
Pathway and disease association enrichment analysis of genes upregulated in low-T cells. Reactome pathway enrichment analysis of genes significantly upregulated in low-T cells relative to high-T cells across tissues. The top 20 pathways ranked by gene count with FDR < 0.05 are shown for (**A**) brain, (**B**) immune cells, and (**C**) lung. DISGENET disease association enrichment analysis of low-T upregulated genes showing the top 20 disease terms ranked by gene count with FDR < 0.05 for (**D**) brain, (**E**) immune cells, and (**F**) lung.

In immune cells, low-T cells show enrichment for cytokine signaling, chemokine receptor interactions, TNF-TNFR pathways, and the TNFR2 non-canonical NF-κB cascade (**Figure 7B, Table 7**). These programs suggest a transcriptionally compressed but functionally persistent immune substate, potentially linked to survival and chronic inflammatory adaptation. In lung, low-T cells show enrichment for metabolic processes, including protein/RNA metabolism, aerobic respiration, respiratory electron transport, ribosome-associated quality control, and stress-responsive pathways such as innate immune signaling and viral response (**Figure 7C, Table 8**). This indicates selective prioritization of metabolic maintenance and stress resilience.

Collectively, across tissues, low-T cells show tissue-specific functional enrichment, yet all share a conserved architecture: selective preservation of essential functional modules and aging-related survival/stress-adaptive pathways.

### Disease associations reinforce biological relevance of low-T programs

We then asked whether the upregulated genes in low-T cells indicate disease-specific associations. DisGeNET analysis revealed tissue-specific disease enrichment consistent with functional programs. In brain, low-T cells were enriched for neuropsychiatric and neurodevelopmental disorders including schizophrenia, bipolar disorder, autism spectrum disorder, intellectual disability, and epilepsy (**Figure 7D, Table 9**). In immune cells, low-T cells showed enrichmed for immune-mediated conditions such as rheumatoid arthritis, hypersensitivity, HIV infection, and celiac disease (**Figure 7E, Table 10**). In lung, low-T cells are upregulated for genes predominantly associated with mitochondrial and metabolic disorders (**Figure 7F, Table 11**).

These associations mirror pathway analysis, reinforcing that low-T states reflect functionally relevant, tissue-specific programs, rather than technical artifacts. Notably, enriched pathways overlap with molecular domains central to aging biology, including synaptic maintenance, chronic inflammation, and mitochondrial function.

### Low-T cells upregulate aging-related genes and vary with sex and age

Having shown aging-related survival/stress adaptive pathways selectively upregulated in low-T cells, we then sought to examine whether low-T cells show presence for human aging genes. Intersection with a curated list of 307 aging-associated human genes^26^ revealed that low-T cells are enriched for aging-relevant genes, suggesting their transcriptional programs may support organ maintenance and tissue resilience rather than senescence, apoptosis, or stress (**Figure 8, Table 12**). To further examine their aging relevance, we quantified the proportion of low-T cells across sex and age. In brain, we found females have higher low-T cell proportions than males (**Figure 9A**). Furthermore, aging decreases low-T abundance in both sexes, with a slightly more pronounced decline in females (**Figure 9A**). In immune cells, low-T cell proportions decreased between 20-year-old and 56-year-old individuals (**Figure 9B**). These results suggest that low-T cell abundance is dynamically modulated by sex and aging, potentially contributing to age-associated changes in tissue homeostasis.

**Figure 8.**
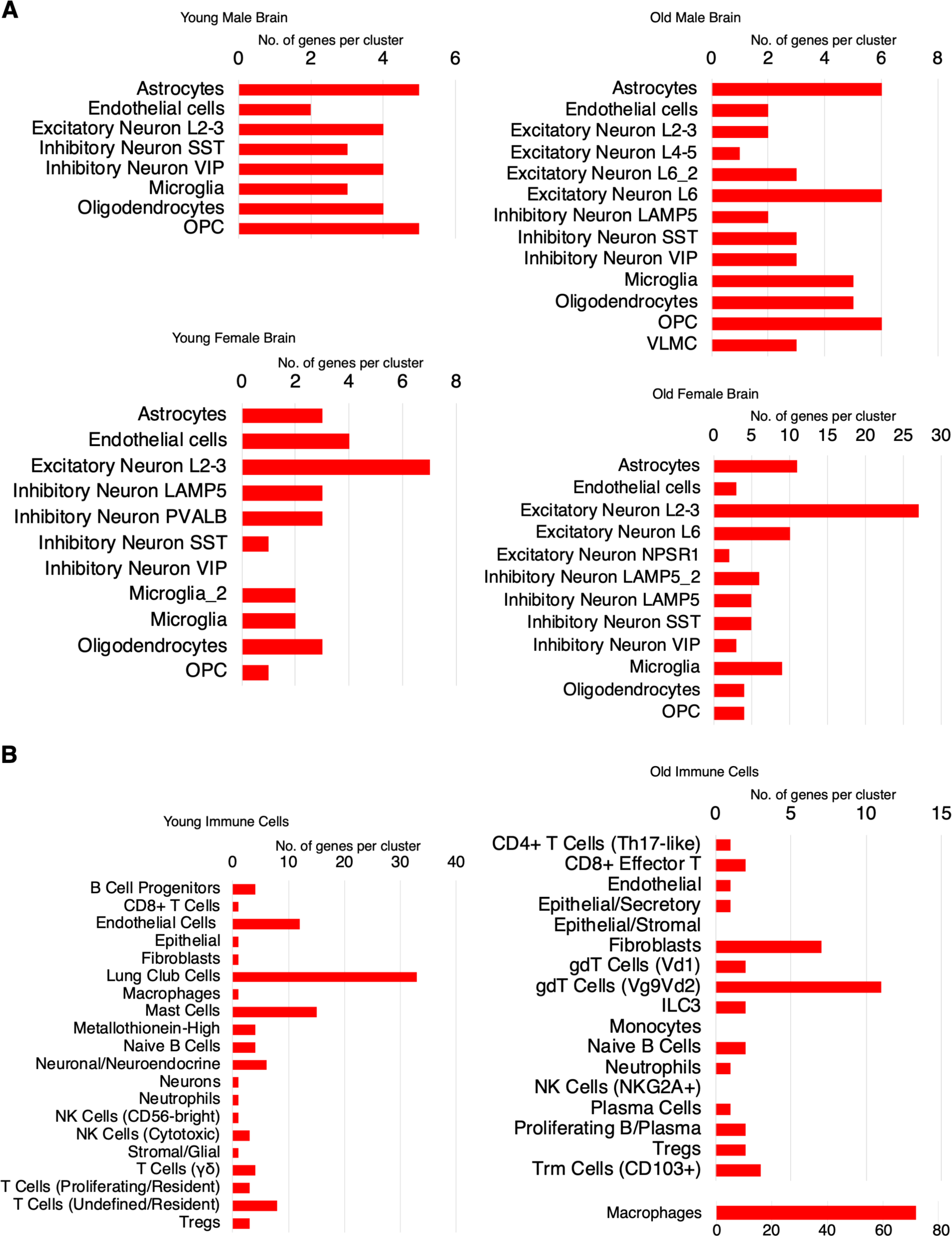
Overlap between low-T upregulated genes and curated human aging-associated genes. Intersection of genes significantly upregulated in low-T cells with a curated list of 307 human aging-associated genes from the GenAge Human Genes database. Intersections are shown for each annotated mature cell type in (**A**) brain and (**B**) immune cell datasets.

**Figure 9.**
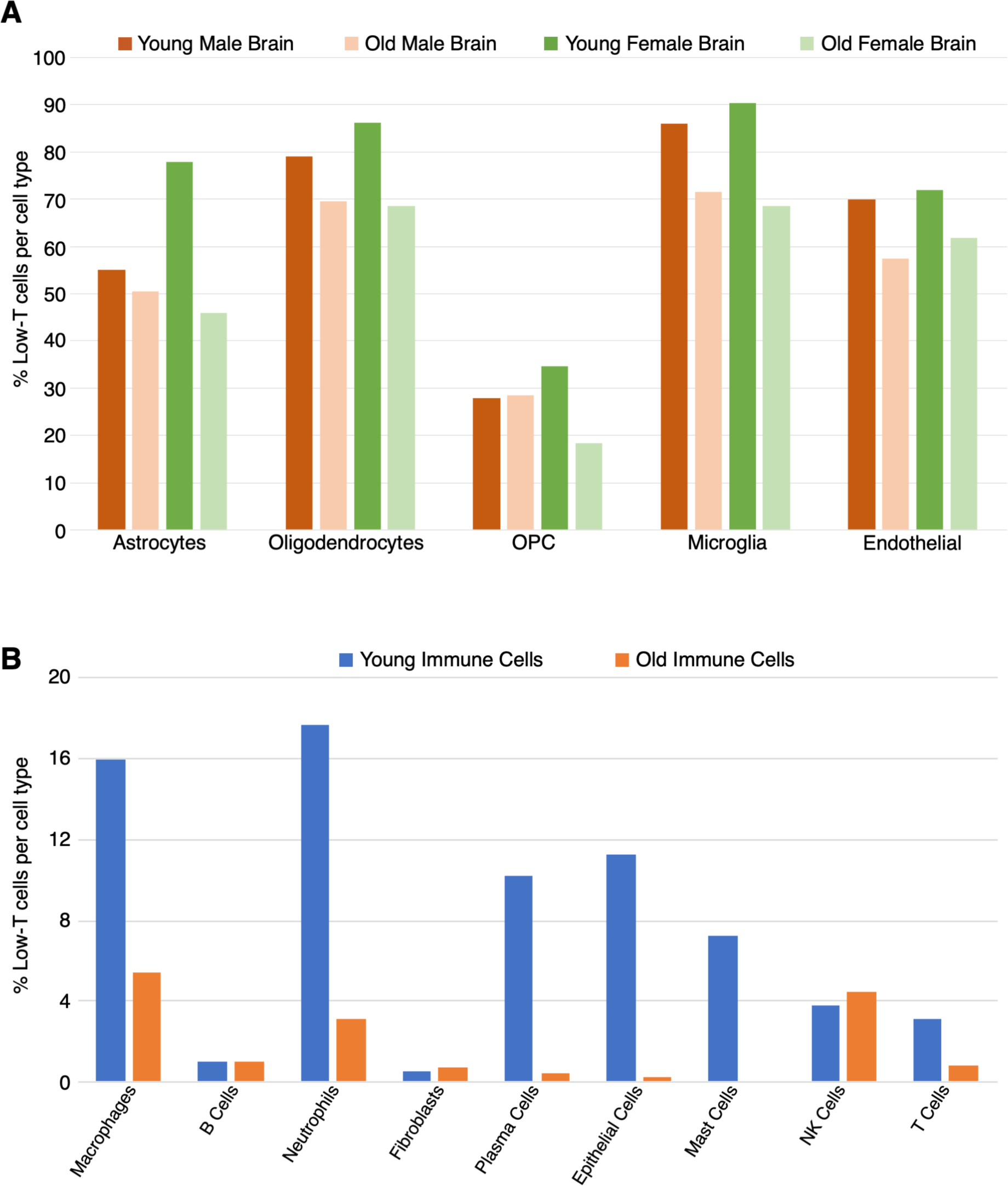
Age-associated differences in low-T cell proportions across brain and immune cell types. Bar plots show the proportion of low-T cells within annotated mature cell types across age groups. (**A**) Human brain datasets stratified by young males, old males, young females, and old females. (**B**) Human immune cell datasets stratified by young and old donors.

## Discussion

Single-cell and single-nucleus transcriptomics have largely operated under the assumption that transcriptionally sparse cells represent technical artifacts, dying cells, or low-quality captures and should be excluded from downstream analysis. Here, we systematically challenge this assumption. We show that low transcriptional complexity cells (“low-T cells”) are widespread across human tissues and constitute a substantial fraction of major mature cell lineages. Across heart, brain, lung, and immune compartments, low-T cells comprised 10-90% of cells within canonical cell types, remained embedded within established clusters rather than forming discrete populations, and exhibited reproducible transcriptional programs distinct from their high-transcriptional (high-T) counterparts. These findings reveal a previously underappreciated dimension of cellular state that is largely invisible to conventional filtering strategies yet appears biologically structured and aging-relevant.

Low-T cells do not reflect stochastic transcript loss or global transcriptional shutdown. Within each lineage, they display selective and coherent gene expression differences, including hundreds of genes consistently upregulated relative to high-T cells of the same annotated type. These differences are qualitative rather than merely reduced transcript abundance, indicating that low-T cells are not diluted versions of high-T cells but represent a distinct molecular configuration. The reproducibility of these signatures across independent datasets, methodologies, tissues, sexes, and ages argues against technical bias as the primary explanation and supports the interpretation that low transcriptional complexity reflects a regulated cellular state embedded within mature populations.

Within each organ, low-T cells share conserved gene programs across diverse mature lineages, whereas overlap across organs is limited. In the brain, multiple glial, neuronal, and vascular populations exhibit shared low-T signatures; similar intra-organ convergence is observed in immune and lung compartments. This pattern suggests that low-T states are governed by tissue-specific regulatory logic rather than representing a universal stress, quiescence, or metabolic shutdown response. Instead, they appear consistent with organ-adapted maintenance programs shaped by shared functional constraints within each tissue environment.

Several observations argue against interpreting low-T cells as dying, stressed, senescent, or apoptotic. Canonical markers of apoptosis, DNA damage, unfolded protein response, oxidative stress, inflammatory signaling, and senescence were not enriched. Low-T cells retained lineage-defining markers and remained embedded within canonical clusters. Moreover, pathway analyses revealed enrichment for programs associated with post-transcriptional regulation, proteostasis, mitochondrial buffering, metabolic resilience, immune restraint, tissue residency, and apoptosis resistance, which are features more consistent with long-lived differentiated cells than with degeneration. If transcriptional sparsity reflected progressive failure, incoherent transcript erosion would be expected; instead, coordinated lineage-consistent programs are observed. Together, these findings support the interpretation that low-T states represent regulated maintenance-oriented configurations rather than artifacts.

A notable feature of low-T cells is their age-associated decline. Across brain and immune datasets, low-T proportions decreased with age, with sex- and cell-type-specific patterns. Low-T states were enriched for genes implicated in human aging and longevity rather than stress pathways. While these data are correlative, they raise the possibility that low-T states reflect maintenance-optimized cellular configurations that diminish during aging. Their reduction may reflect loss of long-lived phenotypes, shifts toward transcriptionally reactive states, or depletion of maintenance-specialized cells. Whether this decline contributes functionally to tissue aging remains to be determined.

Conceptually, our findings suggest that cellular state space includes an orthogonal axis defined by transcriptional output level, with low-T and high-T states coexisting within the same mature lineages. This axis likely reflects differences in transcriptional dynamics, chromatin organization, RNA polymerase engagement, RNA turnover, or post-transcriptional regulation rather than cell fate. High-T states may correspond to transcriptionally adaptive or responsive configurations, whereas low-T states may correspond to stabilized, persistence-oriented modes. It remains unclear whether low-T and high-T cells interact functionally through physical contact, secreted signaling molecules, or extracellular vesicles; however, differential crosstalk between these states represents an important area for future investigation.

These observations have methodological implications. Standard quality-control pipelines frequently exclude cells below fixed gene-count thresholds, potentially removing biologically meaningful states. Such filtering may distort inferred cell-type proportions and obscure aging-associated changes. Notably, all low-T cells analyzed here passed conventional QC metrics and exhibited coherent transcriptional programs, indicating that gene-count thresholds alone are insufficient to distinguish artifacts from regulated states. Stratifying cells by transcriptional complexity within annotated cell types revealed qualitatively distinct configurations overlooked by conventional analyses. Reassessment of QC practices may therefore be warranted, particularly in aging tissues where transcriptional sparsity may characterize long-lived states.

Threshold selection warrants careful consideration. While approximately 1,000 detected genes per cell provided robust stratification across tissues, certain mature cell types exhibited >90% low-T prevalence, suggesting that lineage-specific or adaptive thresholds may better capture biologically meaningful variation. Future studies using finer stratification may clarify the quantitative boundaries that define low-T states.

The tissue-specific programs observed further support the regulated nature of low-T configurations. In the brain, low-T states were enriched for genes associated with electrophysiological stability, synaptic maintenance, RNA processing, and structural integrity. In immune populations, low-T programs emphasized immune restraint, checkpoint signaling, survival pathways, and tissue residency. In lung, enrichment included mitochondrial buffering, proteostasis, antioxidant defense, and apoptosis resistance. Collectively, these patterns are consistent with organ-adapted maintenance strategies that may preserve function while minimizing transcriptional and metabolic burden.

Mechanistically, several possibilities warrant investigation. Low-T states may reflect epigenetically stabilized chromatin landscapes, altered transcriptional bursting kinetics, reduced polymerase engagement, enhanced RNA stability, or coordinated post-transcriptional regulation. Whether these states are reversible or represent stable endpoints within mature lineages remains unknown. Environmental stressors, inflammatory cues, or metabolic conditions may influence transitions between high-T and low-T configurations. Longitudinal lineage tracing and multi-omic integration— including spatial transcriptomics to map whether low-T cells are strategically placed within tissues, chromatin accessibility profiling, proteomics, and live-cell tracking—will be essential to resolve these dynamics.

The relationship between low-T states and aging also suggests potential translational relevance. If low-T proportions track biological aging, they may have biomarker potential, including the development of low-T–restricted transcriptomic or epigenetic clocks. Integration with DNA methylation profiling could determine whether low-T cells exhibit distinct aging signatures. In addition, emerging partial reprogramming strategies in aging biology raise the question of whether interventions broadly targeting tissues preserve, restore, or disrupt low-T states. It is conceivable that maintenance-oriented configurations such as low-T cells may require distinct therapeutic consideration rather than uniform manipulation of entire cell populations.

Several limitations should be acknowledged. Although analyses were restricted to QC-passed cells, technical factors such as capture efficiency, RNA degradation, and nuclear versus cytoplasmic RNA composition may influence gene-count distributions. However, the reproducibility of lineage-embedded low-T programs across datasets, tissues, sexes, and ages, together with the absence of stress signatures, argues against technical artifact as the sole explanation. Our analyses are cross-sectional and observational, limiting causal inference. Functional validation at protein, metabolic, electrophysiological, and in vivo levels remains necessary to define the physiological roles of low-T states.

In summary, low transcriptional complexity cells are widespread, lineage-embedded, tissue-specific cellular configurations enriched for stability and maintenance-associated programs. Their reproducible presence across tissues and decline with age suggest that transcriptional sparsity can encode structured biological information rather than technical noise. Incorporating low-T states into single-cell workflows may refine cellular atlases and provide new insight into mechanisms of tissue maintenance, intercellular organization, and aging biology.

## Competing Interests

Prabhu Mathiyalagan is the CEO and Co-Founder of Benthos Prime Central, TX, USA.

## Online Methods

### Data Acquisition and Pre-processing

Publicly available scRNA-seq or snRNA-seq data files were retrieved from the Gene Expression Omnibus (GEO). Raw count matrices were imported into Seurat 5.3.1 using the function appropriate to the file type deposited in the repository. Initial filtering was performed during object creation to remove cells expressing fewer than 200 genes (min.features = 200) and genes expressed in fewer than 3 cells. The individual datasets representing groupwise were subsequently merged into a single Seurat object as one study group for downstream analysis. This is followed by group wise comparisons.

### Quality Control and Normalization

Quality control was performed to exclude low-quality cells. The percentage of reads mapping to the mitochondrial genome was calculated for each cell (pattern ^MT-). Cells with greater than 10-15% mitochondrial content (percentage determined based on manual inspection of each dataset) were filtered out. Normalization and variance stabilization were performed using SCTransform^27^, during which the mitochondrial percentage was explicitly regressed out (vars.to.regress = “percent.mito”) to minimize the effect of cell stress on clustering.

### Dimensionality Reduction and Batch Correction

Principal Component Analysis (PCA) was calculated on the SCT-transformed data. To correct for batch effects between samples, the Harmony algorithm was applied to the dataset grouping by original identity (orig.ident), utilizing a k-means initialization of 20 and a maximum of 100 iterations^28^.

### Clustering and Visualization

A Shared Nearest Neighbor (SNN) graph was constructed using the Harmony reduction with the first 30 dimensions. Unsupervised clustering was performed using the FindClusters function with a resolution parameter of 0.05 to obtain a reasonable level of granularity. For visualization, a Uniform Manifold Approximation and Projection (UMAP) embedding was generated using the first 20 dimensions of the Harmony reduction. Low transcriptional complexity (low-T, <1000 genes per cell) and high transcriptional complexity (high-T, ≱1000 genes per cell) cellular states were defined based on gene detection per cell and visualized using UMAP embeddings across multiple human tissues.

### Markers and Cell Type Annotation

Cluster markers were identified using the FindAllMarkers function on the “SCT” assay (prepped via PrepSCTFindMarkers). Markers were defined as genes showing positive expression (only.pos = TRUE. The top 20 markers for each cluster, ranked by average log2 fold-change, were extracted and cell annotation was carried out using GPT-4^29^.

### Differential Expression analysis and statistical significance

Differential expression analysis between low-T and high-T cell states within each annotated cell type was carried out and significantly up regulated or down regulated genes were determined using a p-value cut-off <0.05.

### Pathway analysis

Significantly upregulated genes in low-T cell states were used for pathway analysis using Reactome Pathway Database and DISGENET. Only pathways enriched with FDR<0.05 were considered^30,31^.

## Notes

### Summary of Updates

One of the statements lacked citation, which is not corrected and we have added new citations.

## References

1. Kulkarni A, et al. Beyond bulk: a review of single cell transcriptomics methodologies and applications. Curr Opin Biotechnol. 2019 Aug;58:129–136.

2. Inayatullah M et al. Advances in single-cell omics: Transformative applications in basic and clinical research. Curr Opin Cell Biol. 2025 Aug;95:102548.

3. Jovic D et al. Single-cell RNA sequencing technologies and applications: A brief overview. Clin Transl Med. 2022 Mar;12(3):e694.

4. Gulati GS et al. Profiling cell identity and tissue architecture with single-cell and spatial transcriptomics. Nat Rev Mol Cell Biol. 2025 Jan;26(1):11–31.

5. Karlsson M et al. A single-cell type transcriptomics map of human tissues. Sci Adv. 2021 Jul 28;7(31):eabh2169.

6. He S et al. Single-cell transcriptome profiling of an adult human cell atlas of 15 major organs. Genome Biol. 2020 Dec 7;21(1):294.

7. Goldfarbmuren KC et al. Dissecting the cellular specificity of smoking effects and reconstructing lineages in the human airway epithelium. Nat Commun. 2020 May 19;11(1):2485.

8. Chen C et al. Single-cell analysis of adult human heart across healthy and cardiovascular disease patients reveals the cellular landscape underlying SARS-CoV-2 invasion of myocardial tissue through ACE2. J Transl Med. 2023 May 31;21(1):358.

9. Feng W et al. Single-cell transcriptomic analysis identifies murine heart molecular features at embryonic and neonatal stages. Nat Commun. 2022 Dec 27;13(1):7960.

10. Subramanian A et al. Biology-inspired data-driven quality control for scientific discovery in single-cell transcriptomics. Genome Biol. 2022 Dec 27;23(1):267.

11. Yates J et al. Filtering cells with high mitochondrial content depletes viable metabolically altered malignant cell populations in cancer single-cell studies. Genome Biol. 2025 Apr 9;26(1):91.

12. Alvarez-Dominguez JR, Melton DA. Cell maturation: Hallmarks, triggers, and manipulation. Cell. 2022 Jan 20;185(2):235–249.

13. Wigerblad G et al. Single-Cell Analysis Reveals the Range of Transcriptional States of Circulating Human Neutrophils. J Immunol. 2022 Aug 15;209(4):772–782.

14. Sergio Marco Salas et al. Exploration of RNA outside segmented cells in spatial transcriptomics reveals extrasomatic RNA organization. bioRxiv 2025.12.07.692889; doi: 10.64898/2025.12.07.692889.

15. Ibáñez-Molero S et al. Tumour-reactive heterotypic CD8 T cell clusters from clinical samples. Nature. 2026 Jan;649(8096):467–476.

16. Ji S et al. Cellular rejuvenation: molecular mechanisms and potential therapeutic interventions for diseases. Signal Transduct Target Ther. 2023 Mar 14;8(1):116.

17. Yun MH. Changes in Regenerative Capacity through Lifespan. Int J Mol Sci. 2015 Oct 23;16(10):25392–432.

18. Liu F et al. Single-cell mitochondrial sequencing reveals low-frequency mitochondrial mutations in naturally aging mice. Aging Cell. 2024 Sep;23(9):e14242.

19. Xu X et al. Mitochondria in oxidative stress, inflammation and aging: from mechanisms to therapeutic advances. Signal Transduct Target Ther. 2025 Jun 11;10(1):190.

20. Palmer AK, Jensen MD. Metabolic changes in aging humans: current evidence and therapeutic strategies. J Clin Invest. 2022 Aug 15;132(16):e158451.

21. Singh A et al. Aging and Inflammation. Cold Spring Harb Perspect Med. 2024 Jun 3;14(6):a041197.

22. Debès C et al. Ageing-associated changes in transcriptional elongation influence longevity. Nature. 2023 Apr;616(7958):814–821.

23. Santra M et al. Proteostasis collapse is a driver of cell aging and death. Proc Natl Acad Sci U S A. 2019 Oct 29;116(44):22173–22178. doi: 10.1073/pnas.1906592116.

24. Jeffries AM et al. Single-cell transcriptomic and genomic changes in the ageing human brain. Nature. 2025 Oct;646(8085):657–666.

25. Zhang Z et al. A panoramic view of cell population dynamics in mammalian aging. Science. 2025 Jan 17;387(6731):eadn3949.

26. de Magalhães JP et al. Human Ageing Genomic Resources: updates on key databases in ageing research. Nucleic Acids Res. 2024 Jan 5;52(D1):D900-D908.

27. Hafemeister C, Satija R. Normalization and variance stabilization of single-cell RNA-seq data using regularized negative binomial regression. Genome Biol. 2019 Dec 23;20(1):296.

28. Korsunsky I et al. Fast, sensitive and accurate integration of single-cell data with Harmony. Nat Methods. 2019 Dec;16(12):1289–1296.

29. Hou W, Ji Z. Assessing GPT-4 for cell type annotation in single-cell RNA-seq analysis. Nat Methods. 2024 Aug;21(8):1462–1465. doi: 10.1038/s41592-024-02235-4.

30. Milacic M et al. The Reactome Pathway Knowledgebase 2024. Nucleic Acids Res. 2024 Jan 5;52(D1):D672-D678.

31. Janet Piñero et al. DISGENET: Accelerating Data-Driven Discovery in Disease Genomics and Therapeutic Development. bioRxiv 2026.01.05.697749; doi: 10.64898/2026.01.05.697749.

